# Interaction studies of Gut metabolite; Trimethylene amine Oxide with Bovine Serum Albumin through Spectroscopic, DFT and Molecular Docking Approach

**DOI:** 10.1101/2023.04.06.535846

**Authors:** Awadhesh Kumar Verma, Payal Gulati, GBVS Lakshmi, Pratima R Solanki, Anil Kumar

## Abstract

Trimethyleneamine N-oxide (TMAO); a gut microbiota derived metabolite has been involved in human health and diseases. It is enhanced by insulin resistivity and linked with various metabolic syndromes in human being such as renal, neuro-degenerative, and cardiovascular diseases. The primary mechanism through which TMAOs promotes disease is not clear yet. TMAO with MW= 75.11 g/mol is a small biomolecule hence, it becomes crucial to develop the conjugate of TMAO with BSA for aptamer synthesis. The binding interactions among TMAO and BSA were investigated using spectroscopic methods like UV-Vis, photoluminescence, Fourier transform infrared and circular dichroism. Hydrophilicity/Hydrophobicity of the conjugate was monitored by using contact angle (Ɵ) measurement. Sodium dodecyl sulphate polyacryl amide gel electrophoresis (SDS-PAGE) confirmed the different ratio of conjugate formation with the help of band size. This interaction study reveals that TMAO bind with BSA on two sites and with high affinity on one site. Docking studies also showed TMAO is involved in non-covalent interaction with bovine serum albumin forming stable docking complex with binding score of –3.6 kcal/mol obtained from the docking simulation. TMAO is involved in interaction with BSA via amino acid residues forming the stable docking complex through hydrogen bond and electrostatic interaction. This kind of interaction study may be helpful in making strategies to break the conjugation between serum albumin and uremic toxin and pave the way for the treatment for CKD and other diseases wherein TMAO is implicated. Also, conjugation of TMAO and BSA studied here may also serve as premise to develop aptamers for the detection of TMAO in the body fluids.

## 1. Introduction

Trimethylamine N-oxide (TMAO); is very small and colorless organic amine oxide. It is one of the most pronounced metabolite derived from some gut microbes which have some adverse effect on human health causing so many biological problems like DNA damage, oxidative stress, protein misfolding, inflammation, cardiovascular disorder like Atherosclerosis Cardiovascular Disease (ACVD), heart attack, cerebrovascular, Alzheimer’s, metabolic syndrome, Diabetes mellitus type II, Insulin resistance, chronic kidney diseases and colo-rectal cancer (CRC) [1]–[4]. It gets accumulated inside the tissues of sea animals in very high concentrations. It acts as safeguard against protein destabilization effect of urea[5]. Plasma TMAO level is regulated by several factors like gut microbial flora, diet and flavin mono-oxygenase activity inside the lever. The atherogenicity of TMAO is accredited to the alteration of bile acid and cholesterol metabolism, inflammatory pathways activation and by the promotion of foam-cell i.e., lipid laden macrophage formation. Levels of TMAO increases with decrease in kidney function and is related to the mortality of patients suffering from chronic kidney diseases.

TMAO is originated from trimethylamine (TMA) that is metabolized from several precursors present in dietary food like choline, betaine, and carnitine metabolized through gut microbes shown in Fig. 1. Gut bacteria metabolize the choline and choline containing compounds ingested directly from dietary food giving rise to intermediate metabolite trimethylamine (TMA) that finally converted in to TMAO in liver. Flavin-dependent monooxygenase (FMO1 and FMO3) converts TMA into TMAO inside the liver. Activities of these FMO’s is related to their genetic flippancy. TMAO production is directly related to gluconeogenesis and glucose transport occurs in liver that leads to increased rate of susceptibility towards insulin resistance. So, we can suggest that TMAO may be considered as causal link between insulin resistance development and cancer. So, we can reduce the TMAO formation either via limiting the consumption of precursors of TMAO or by bacterial growth inhibition that catalyzes the formation of TMAO and hence the risk of diabetes or cancer may be reduced.

**Fig. 1:**
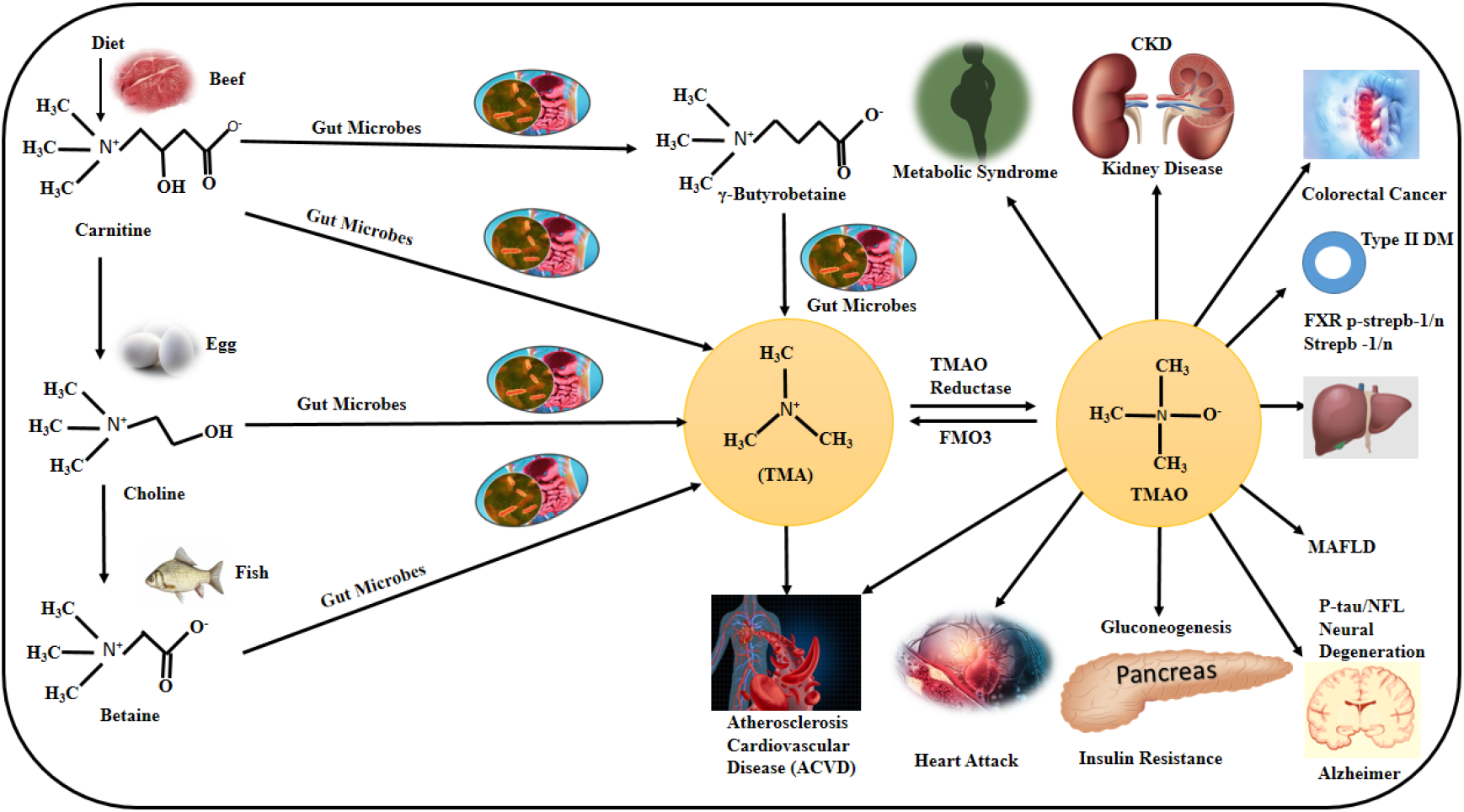
Schematics of molecular pathway for formation of TMAO from various precursor obtained from ingested foods [6].

TMAO has the capability to alter the structure as well as activities of so many compounds of biological importance. TMAO acts as protein and nucleic acid stabilizer in folded state[7]. Several thermodynamical studies have confirmed that TMAO prevent the protein form it’s denaturation through heat, pressure and protein denaturants[8]. Various mechanisms have been put forward to explain the protein folding propensity of TMAO [9], [10]. It has been hypothesized that TMAO can affect the protein stability by altering the structure of water as well as strength of hydrogen bonding. Numerous studies have signified the robust interactions of hydrophilic groups present in TMAO with the water molecules for homolytic activity of TMAO. The insinuation from these kinds of findings it can be said that TMAO can stabilizes the proteins by altering the polarity of protein and water activity. It might affect the equilibrium between unfolded and folded state of proteins indirectly [11], [12]. However, various molecular dynamics simulation study has negated the alteration of water structure in aqueous solutions of TMAO[13]. As the mechanism of specific of TMAO with Bovine serum albumin (BSA) and the effect of TMAO on protein confirmation are still not clear. So efforts have been made to study these aspects in details through this study [14]. Serum albumin (SA) is the superabundant proteinaceous component of blood plasma that promotes the transportation as well as disposition of several exogenous as well as endogenous ligands including metal ions, fatty acids and steroids[14]. It is also responsible to maintain the pH of blood and contribute the colloidal osmotic blood pressure. Fatty acid and dye absorption, their distribution and metabolism along with property of excretion attributes as well as toxicity including stability can be remarkably affected by their binding to the serum albumins. In addition, it is evident that conformational change in serum albumin is prompted by interaction with metabolites, drugs molecules and dyes having lower molecular weight, that significantly alter the albumin secondary and tertiary structure[14]. Therefore, conjugation of metabolites, drugs molecules and dyes with serum albumin is very important in the field of biomedical research. If we consider the circulating baseline TMAO levels in male and female it does not make significant difference as in females it is 2.85 ± 1.64 μmol/l, p = 0.89 while in males it is 2.82 ± 1.60 μmol/l [15]. Since the generation of TMAO is extremely depending on gut microbiota and composition of gut that is associated to insulin resistivity and hence type II diabetes. Therefore, drugs molecule which alter the microbial development may be considered as potential preventive therapeutic target for both insulin resistivity as well as cancer risk.

In the several research studies it has been found that TMAO can be considered as the best prognostic and diagnostic biomarker for cardiovascular, diabetes, colorectal cancer (CRC), and other diseases. Also, it has been established that TMAO can act as the early marker for the detection of atherosclerosis. TMAO has unique chemical properties which makes it an electron acceptor, an osmolyte and stabilizer of protein and hence involved in osmotic, oxidative and hydrostatic stress like situations. Also regulates the activity of various hormones and enzymes[2]. Molecular docking or simply docking is an *in silico* approach, which endeavor to mimic the interaction of ligand molecules to any biomolecules like nucleic acid and protein[16]–[20]. It anticipates the optimized binding conformations. The orientations of molecules binding with one another in 3-D space along with evaluation of binding affinity, scoring function and binding energy which may represent the strength of binding, 3-D structure along with complex stability[21]–[23]. It confer the comprehensive knowledge about the fitting of ligand in to the active site, proper orientation of ligand molecule and most stable confirmation in the binding pocket including de novo designing and ligand optimization[24], [25]. It provides conglomerate analyses depending on the diverse range of scoring methodology and particular selection criteria for binding[26]. It mainly uses the proposition of reciprocity of biomolecules in which the structure gets interacted with one another like hands in a gloves, where both shape of structure as well as physicochemical properties vouchsafe best fitting situation among biomolecules.[27] It forecast position and orientation of the ligand in specific binding pocket. It is regarded as an essential constituent in the current modern drug discovery edge [28]. In current era, almost all pharmaceutical and biotech and companies are utilizing this methods most routinely and extensively for diverse area of applications[29].

### 2. Experimental Section

#### 2.1 Reagents and Buffer

Bovine Serum Albumin (BSA) in lyophilized form, Tetramethyl ethylene diamine, Sodium dodecyl sulfate (SDS), Acryl amide, N, N methyl acrylamide, Bromo phenol blue and Trimethylamine N-Oxide (TMAO) with high purity were procured from Sigma Aldrich. Dialysis membrane (Snake Skin™ Dialyis Tubing, 10K MWtO, 16mm) was procured from Thermo Scientific. Glacial acetic acid was procured from TM Media. Sodium acetate was procured from Merck. Tris base was procured from G Bioscience. Ammonium per sulfate and glycine was procured from HiMedia. Comassie brilliant blue dye methanol and ethanol were procured from SRL Ranbaxy. 0.2-micron filter was procured from MDI and 2ml syringe was procured form HMD. MilliQ (18.2 MΩ·cm @ 25 °C) from the Millipore water purification system was used for preparation of all the solutions. Acetate buffer at pH=4.3 was used for TMAO-BSA conjugate preparation.

#### 2.2 Apparatus and Instrumentation

The UV–vis absorption spectra of BSA were taken in the absence and presence of TMAO using UV-2600, UV-VIS Spectro-photometer SHIMADZU (Shimadzu Corporation in Japan), through 1.0 cm quartz cells keeping slit width 5 nm and 600 nm/minute scan speed. The spectra were recorded in wavelength range between 200-600 nm. Fluorescence was measured with the help of Fluorescence spectrophotometer (Cary Eclipse, Model G9800A, Agilent Technology,) with fixed excitation as well as emission slit width (5 nm) and 600 nm/minute scan speed. A quartz cuvette (3.5 ml) with path length 10 mm was used. To determine the functional groups changes, residing on the surfaces of bio-conjugates; FTIR spectrophotometer were used in ATR mode (PerkinElmer Spectrum 1). The spectra were obtained in between range of 4000 - 400 wavenumber, with scan speed of 32. The contact angle measurement was done using drop shape analyzer (KRUSS, Germany) to check the hydrophobicity/hydrophilicity of the TMAO-BSA conjugate on ELISA plate. CD spectroscopic study was performed monitor the effect of TMAO on secondary and tertiary structure of BSA in acetate buffer solution. CD spectra were obtained in the UV region using JASCO-1500 CD spectropolarimeter associated with a Peltier temperature controller (PTC-517). Prior to the measurement, instrument was calibrated routinely with D-10 camphor-sulfonic acid under N_2_ purging in lamp compartment, optics and sample chamber in a ratio of 1:3:1 to remove oxygen content and create the inert atmosphere. CD spectra i.e., ellipticity of the acetate buffer solution was recorded as a reference/background in the UV range from 190-250 nm using a cell (path length 1 mm) at room temperature (25°C), 1 nm bandwidth. The average of three measurements were displayed as final spectra with mean time 1 nm/sec. SDS-PAG Electrophoresis was carried out on Mini-PROTEAN®3 Cell apparatus (Bio-Rad). A 12% resolving gel (7 cm in height) was casted in the gel casted followed by topping with 5% stacking gel (1cm in height). The gel cassette was taken out from casting stand and then placed independently in the electrode’s assembly. The clamp stand is fixed to the electrode assembly. A 1X running buffer was added to the opening of the casting frame to fill the wells of the gels. 30 μL samples of different molar ratio of TMAO-BSA conjugate were loaded in different wells with 9 μL of 4X loading dye was added to the wells. Initially the gel was supplied with 60 V to pass the bands from stacking gel and thereafter, voltage was increased to 80 V to run the gel.

#### 2.3 Conjugate Preparation

TMAO (in acetate buffer, PH=4.3) was conjugated with BSA (acetate buffer, pH=4.3) in different molar ratio’s including 1:1, 1:5, 1:10, 1:25, 1:50, 1;75, 1:100. A 3.5 mM of TMAO was mixed with 2μM of BSA and allowed to react for 2 h at room temperature and for 72 h at 4°C. At the end of the reaction, dialysis was performed for 72 h with change of acetate buffer (pH=4.3) after every 12 h.

#### 2.4 Computational Details

We have theoretically modelled 3D structure of gut metabolite TMAO and went for the optimization and energy minimization of the molecule using Marvin Sketch, Chemdraw, Chem3DPro and Gaussian09 software. BSA structure (Source organism: *Bos Taurus*, resolution: 2.70 Å and having 583 amino acid residues) was retrieved from Protein Data Bank having RCSB PDB ID: 3V03. The structure was cleaned of crystallographic water molecules and other crystalizing agents. Docking was performed to find the binding affinity and binding energy of TMAO-BSA conjugate using software AutoDock 4.2., VMD and PyMol software have been used for visualization and identification of the amino acid residues in active site.

## 3. Results and Discussion

Interaction between metabolite (TMAO) and BSA is studied using following spectroscopic techniques.

### 3.1 UV-Visible spectroscopy and PL study

UV–visible absorbance spectra of TMAO, BSA and TMAO-BSA (deducting TMAO absorbance spectrum in acetate buffer, pH 4.3) solutions measured separately to ensure the mechanism of quenching. Fig. 2 (a) shows that absorbance intensity of BSA around 280 nm decreases with the stepwise addition of TMAO solution in different concentrations. This suggests that quenching in absorbance intensity of BSA occurred mainly due to TMAO-BSA conjugate formation[30]. The weak peak obtained at 280 nm resulted from the aromatic amino acids (Try, Tyr and Phe) in protein structure. Proteins absorb UV radiation proportional to the amino acid content present in it. Therefore, it is possible to quantify proteins depending upon the absorption intensity. Aromatic amino acids like tryptophan and tyrosine shows absorbance of UV light at 280 nm which is due to presence of aromatic ring structure in their side chain R group. Delocalization of π electrons takes place within the aromatic ring, that is responsible for the absorption of light by aromatic amino acid residues.

**Fig. 2:**
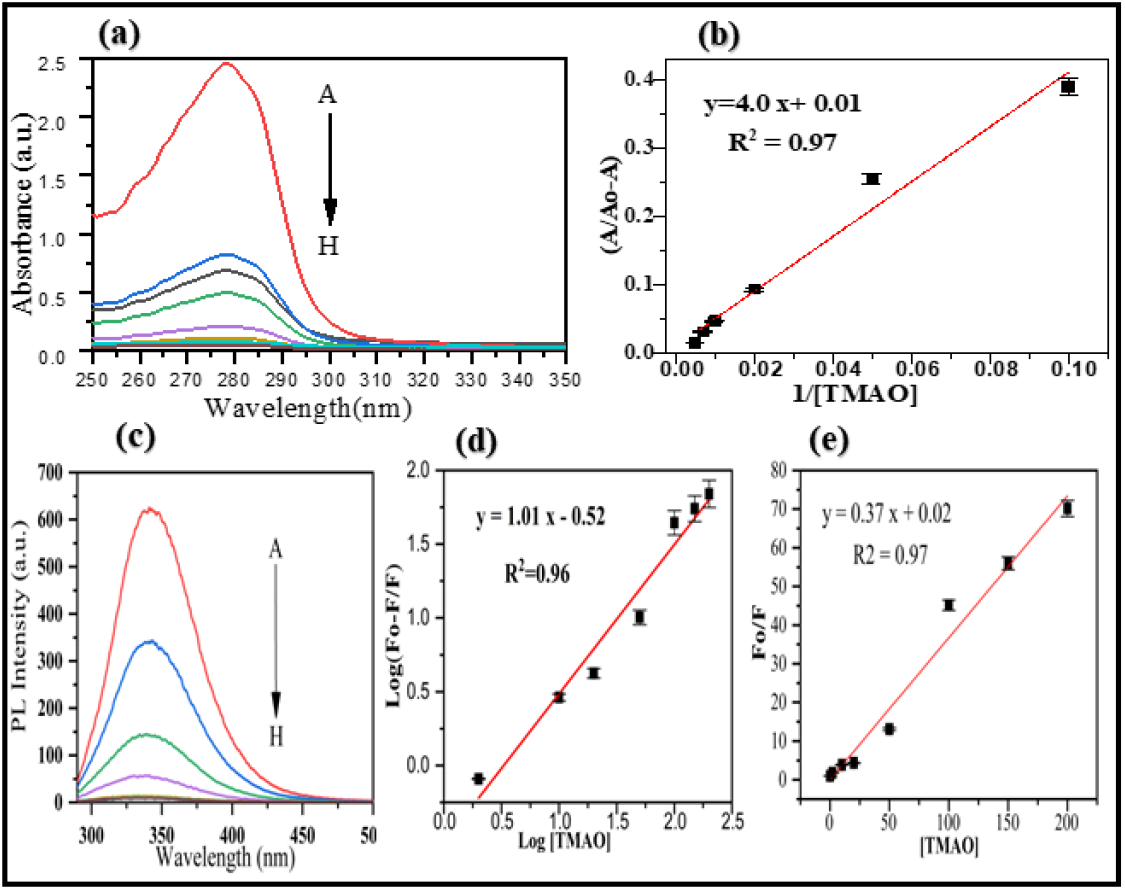
(a) UV-Vis absorbance spectra recorded for various concentration of TMAO in fix concentration of BSA, (b) Benesi-Hildebrand plot UV-Visible plot (A/A_0_-A v/s 1/ [TMAO]) to calculate the total number of binding sites located on the BSA structure to associate with the TMAO. (c) PL spectra of BSA in the absence and presence of TMAO at different concentration (d) Stern–Volmer (SV) plot (e) Fo/F v/s [TMAO].

Benesi-Hildebrand plot (Fig. 2 (b)) was derived from the UV-Visible plot (A/A_0_-A v/s 1/ [TMAO]) to calculate the total number of binding sites located on the BSA structure to associate with the metabolites. TMAO Slope of this plot gives the value of number of binding sites that is 4. The obtained values are in well agreement with the computational data.

The phenomenon of quenching of fluorescence is decrement of quantum yield of fluorophore fluorescence induced by various types of interactions at molecular level, like ground-state conjugate formation, excited-state reactions, quenching due to collision and energy transfer[30]– [34]. By measuring intrinsic PL intensity of BSA protein before and after adding TMAO, there are some changes in microenvironment in proximity of fluorophore molecules. (Fig. 2 (c)) depicts the PL spectrum of BSA in the presence of different concentrations of TMAO at room temperature. When various amount TMAO was mixed with BSA solution which is fixed in concentration, the PL intensity of BSA around 280 nm observed to be decreasing in regular pattern but λmax (maximum emission intensity) remained unchanged did not change to either longer or shorter wavelength.

This signifies that TMAO might have interacted with BSA and quenched its intrinsic PL intensity, but here there was not any changes observed in local dielectric micro-environment of BSA. (Fig. 2 (d & e)) shows to Stern–Volmer (SV) plot at room temperature. Graph shows that for the given investigated range of concentrations, the SV plot shows a good linearity. The Stern-Volmer constant (Ksv) may be obtained from both static as well as dynamic components. SV equation related to static quenching is formulated by taking consideration of conjugate formation between fluorophore molecule (F) and quencher molecule (Q) as reversible reaction along with association constant (K). But where there is dominancy of static quenching, the Ksv may be considered to be equal to association constant (K) between fluorophore and quencher molecule[35]–[37]. The intensity of fluorescence is considered as directly proportional to the concentration of fluorophore. The initial concentration of fluorophore [F_o_], is equivalent to addition of the free fluorophore concentration [F], and nonfluorescent conjugate [FQ]. This gives linearity for Stern-Volmer equation for static quenching.

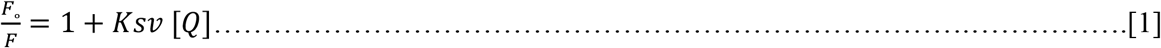

where F_0_ is the PL intensity in absence of quencher molecule, while F in the presence of quencher molecule. K_SV_ is SV quenching constant and [Q] is the quencher concentration. Here results show an excellent linear correlation between (F_0_/F_1_) and [TMAO]. The coefficient K_SV_ is equal to the slope of this line; in this case, the value is 0.37.

From the above discussion it clear that here static quenching of BSA has been occurred induced by TMAO. The data of PL quenching of BSA protein was analyzed to find the several binding factors. The total number of binding pocket (n) and binding constant (K_b_) may be calculated according to below equation [38]

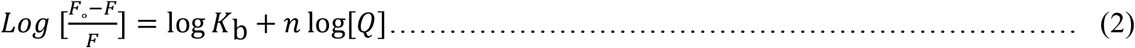

Here F_o_ and F are fluorescent intensities in absence and presence of quencher molecule in steady state. From the Eq. (2), values of n and K_b_ at room temperature were obtained to be 1.01 and 0.28 respectively. This implies that TMAO molecule is strongly bound to BSA. Also, here there is one independent class of binding pocket for TMAO molecule towards BSA. The linear-coefficient (R) is 0.96 which indicates that the underlying assumptions of derivation for equation (2) was satisfactory.

To determine the interaction force between TMAO and BSA protein, the signs as well as magnitudes of the thermodynamic parameter (ΔG) are accountable for the main interaction forces involved in binding process. The force of interaction between ligands and bio-macromolecules includes multiple hydrogen bonds, hydrophobic interaction, electrostatic interactions and van der Waals forces etc. Change in free energy (ΔG) was further estimated from the equation mentioned below:

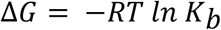

where K_b_ and R represents binding constant and gas constant (8.314 J/Mol/K) respectively. The negative value of ΔG indicates the spontaneity of the reaction between metabolite and BSA. The negative value of ΔG (−3.15 KJ) revealed that binding process is spontaneous.

### 3.3 CD study

To observe the secondary structural change in BSA protein after interacting with ligand molecule, circular dichroism spectroscopy technique was used [39]. Fig. 3 shows the CD spectrum of the BSA interaction with different concentration of the TMAO in acetate buffer pH=4.3 at room temperature. Fig. 3 clearly shows that BSA is exhibiting two negative peaks. One at 208 and other at 222 nm in ultraviolet zone, which is characteristics of typical α-helix structure of BSA protein. Both peaks 208 as well as 222 nm both contributing towards n→π* transition for the given peptide bonds of α-helices. As the concentration of TMAO is increased here, the intensity curves of given decreases in regular pattern from A–H (Fig. 3, band intensity curves A–H). The CD spectra of BSA protein were taken in the presence as well as in absence of TMAO that are showing similar shape of BSA indicating that BSA structure is still predominating the α-helices.

**Fig. 3:**
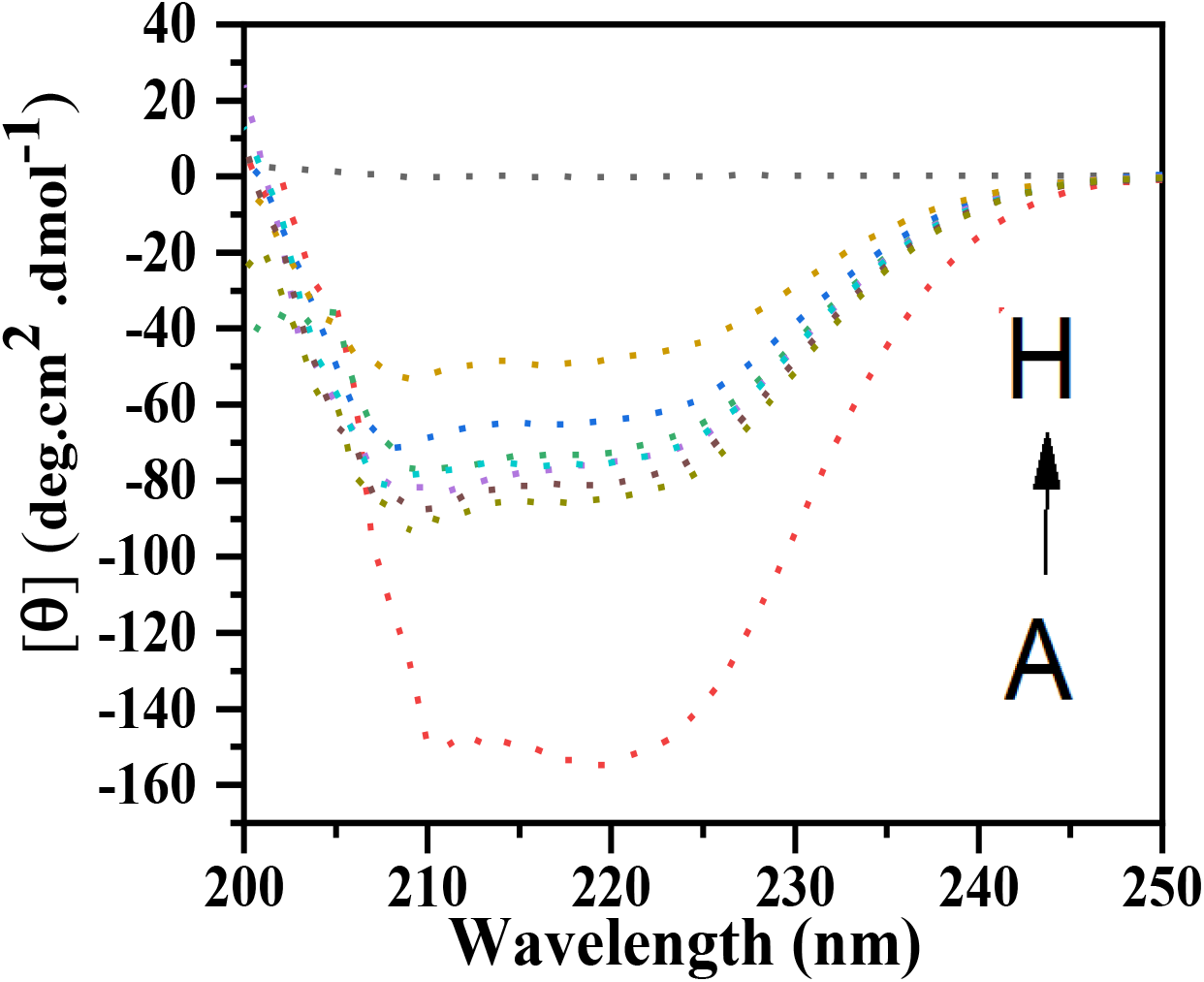
CD spectra of BSA recorded in the presence and absence of TMAO at various concentration.

### 3.4 FTIR

To confirm the interaction between TMAO-BSA conjugate, the FTIR spectral analysis was used. The compositions of pure BSA, TMAO dissolved in acetate buffer pH = 4.3, and TMAO-BSA conjugate were tested separately. Fig. 4 shows the comparison of the FTIR spectra for pure BSA, TMAO in acetate buffer and TMAO-BSA conjugate [40], [41].

**Table 1.** Transmission peaks in FTIR and their respective assignment in BSA, TMAO and TMAO-BSA conjugate.

| Wavenumber<br>(cm<sup>-1</sup>) | Assigned To | TMAO | BSA | TMAO+BSA |
| --- | --- | --- | --- | --- |
| 956 | C-C bond out of the plane<br>deformation | √ | √ | √ |
| 1075 | C-OH stretch | √ (due to<br>acetate buffer) | -- | √ |
| 1146 | C-N bending | √ | √ | Shifted to 1139 cm<sup>-1</sup> |
| 1243 | N-O in N-Oxides | √ |  | Shifted to 1253 cm<sup>-1</sup> |
| 1344 | N-H and C-H in-plane deformation | √ | √ | √ |
| 1400 | C-N stretching vibration | √ |  |  |
| 1465 | CH<sub>3</sub> deformation stretch | -- | √ | -- |
| 1562 | N-O in aliphatic nitro compounds | √ | √ | Shifted to 1548 cm<sup>-1</sup> |
| 1640 | C=O stretch | √ (due to acetate buffer) | √ | √ |
| 1656 | NH<sub>2</sub> stretch | -- | √ | √ |
| 1735 | C=O stretch | -- | √ | √ |
| 1964 | C=C Antisymmetric Stretch | √ | √ | √ |
| 2030 | *N – H* deformation stretch | √ | √ | √ |
| 2150 | N-C Stretch | √ | √ | √ |

**Fig. 4:**
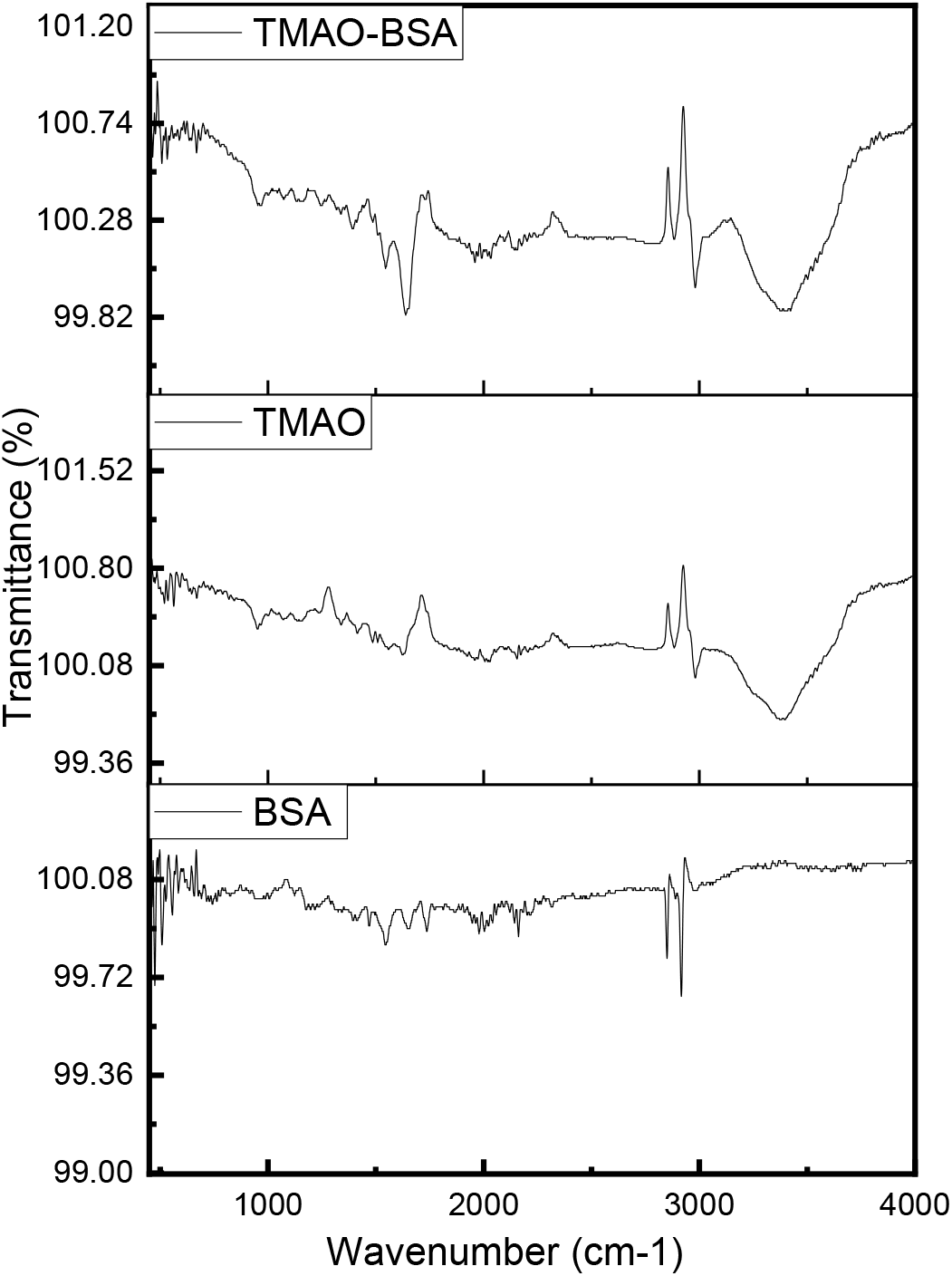
FTIR spectra of bovine serum albumin recorded in the presence and absence of TMAO at various concentration.

There are few peaks in the fingerprint region from 900 to 1650 cm^-1^ which were present in all three spectra corresponding to C-OH, C-N and COO-groups that are due to both BSA and TMAO. The CH stretch in aliphatic compounds appeared between 2800-3000 cm^-1^ which were present in all three spectra. The broad peaks appeared above 3000 cm^-1^ correspond to OH and NH stretch vibrations. From the above table it is also observed that there is a shift in few peaks at 1146, 1243 and 1562 cm^-1^ in conjugation spectrum indicating the interaction between both the components. Thus, FTIR spectrum confirmed the formation of conjugation between TMAO and BSA.

### 3.5 Contact Angle

Measurement of contact angle of BSA, TMAO and conjugates (BSA: TMAO) was carried out on ELISA plate. The contact angle of plate is 77 ° with ionic liquid (IL) which acetate buffer containing ions in it. The stronger the interaction of the ionic liquid ions among themselves i.e., stronger the hydrogen bonding of IL, lesser is its interaction with the surface. Thus, making a greater bond angle with the solid-liquid interface. Further, the contact angle formed by BSA molecules with ELISA plate is 82.95° which means this protein is hydrophobic in nature and is involved in hydrophobic-hydrophobic interaction with the plate surface. Subsequently, Ɵ value for TMAO was also check which is less in comparison with BSA. This suggests the TMAO molecules form stronger bond with plate surface in comparison to the TMAO in acetate buffer. Thereafter, when TMAO was allowed to form conjugate with BSA molecules in different ratios from 1:1 to 1:100; the Ɵ value continuously decreased with the increasing ratio, respectively. TMAO is amphiphilic molecule in nature as it has both hydrophilic and hydrophobic moieties [42] shown in Fig.5.

**Fig. 5:**
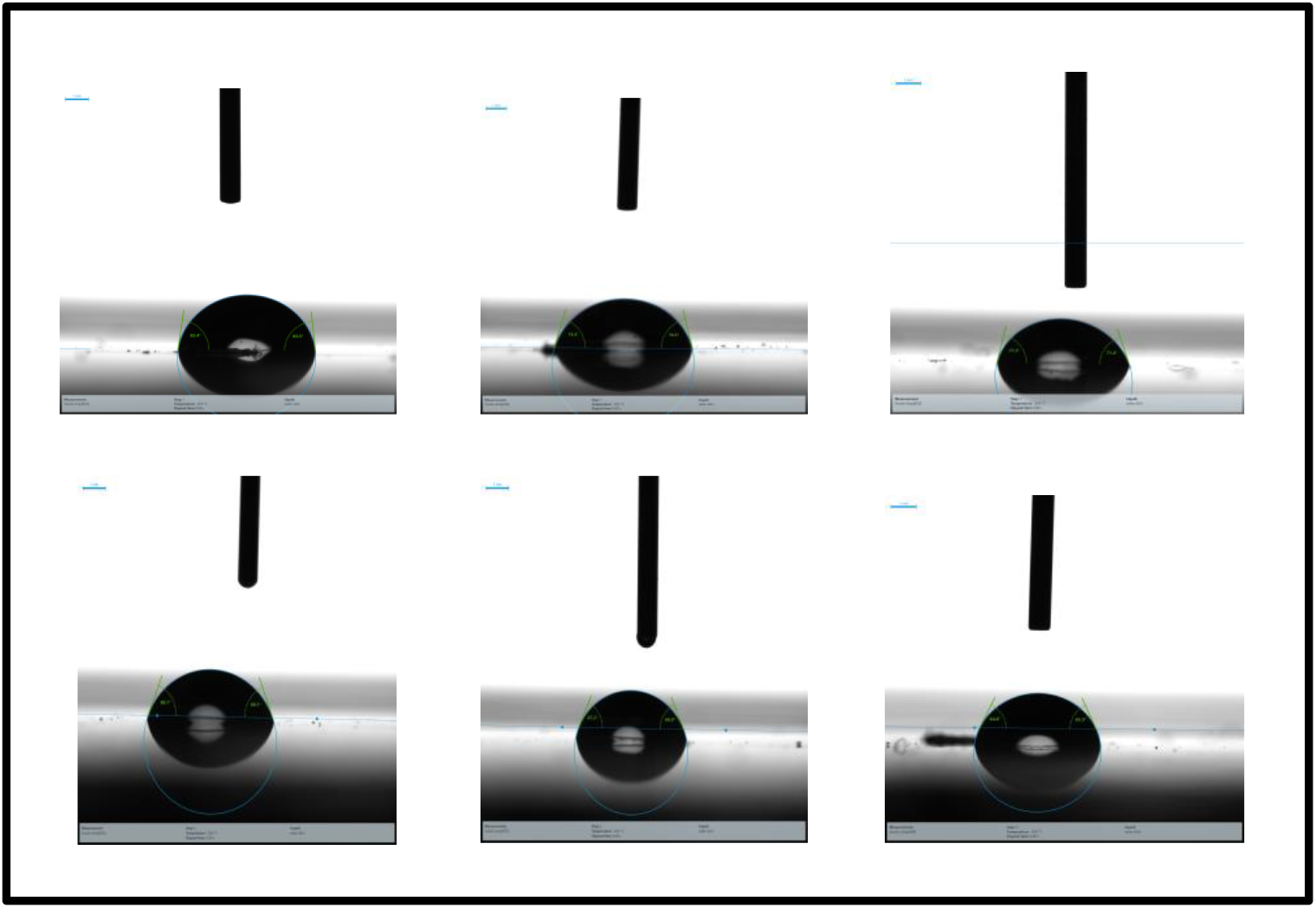
Contact angle of bovine serum albumin and of TMAO-BSA conjugate at various concentration of TMAO.

### 3.6 SDS-PAGE

The prepared conjugates in different ratios including BSA and TMAO were identified using SDS-PAGE technique. The molecular weight of BSA-metabolite was different from BSA as compared to BSA sample as control. The molecular weight of all the seven conjugates were decreased, as shown in Fig. 6. This indicates that the metabolite was successfully conjugated to the BSA carrier proteins. First lane represents the ladder which on resolving shows different molecular weight proteins and helps in identifying the size of the unknown loaded samples. Second lane shows the native BSA and rest lanes so on show conjugates ranging from 1:1 to 1:100. The results obtained suggest that BSA with weight 66 KDa when coupled with different concentration and ratio of metabolite were depicted in the attained band intensity. On increasing the metabolite ratio and kept the BSA in fixed amount, the band intensity diminished significantly, and these results were in good agreement with FL intensity. As SDS-PAGE identifies only proteins and therefore, the band size was reduced as represented in Fig. 6.

**Fig. 6:**
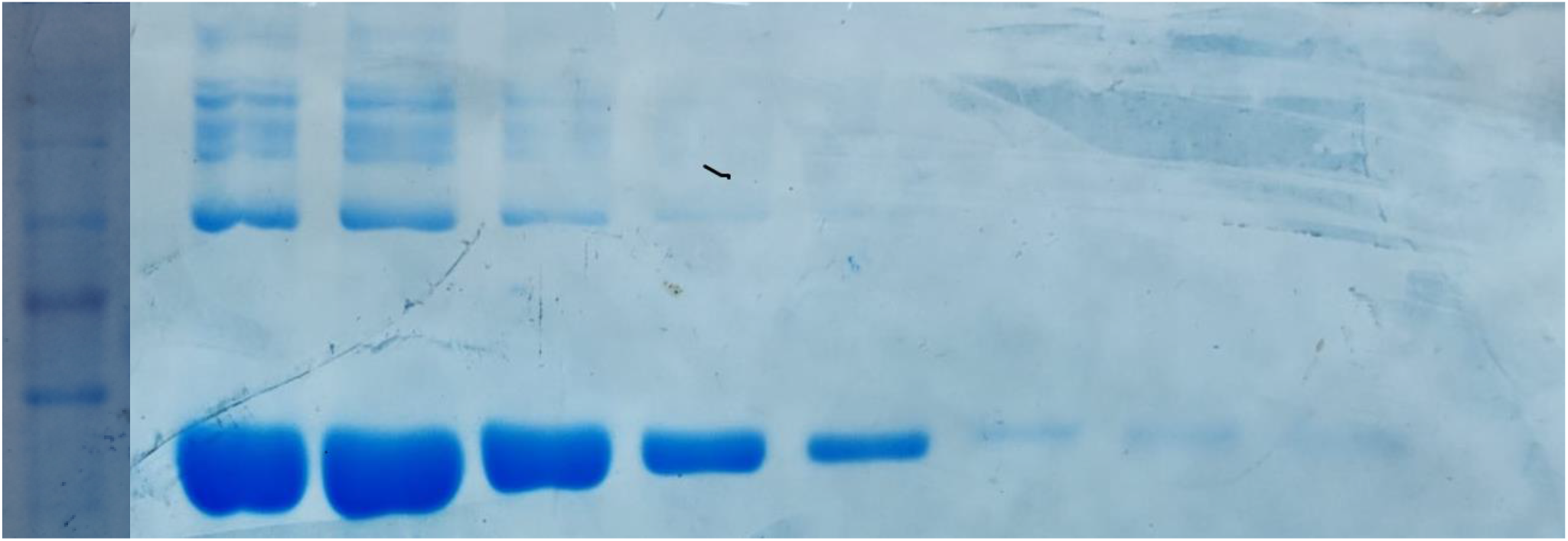
SDS PAGE of BSA and TMAO-BSA conjugate at various concentration of TMAO.

### 3.7 Docking Results

#### 3.7.1 Theoretical Modeling of TMAO

3D structure of TMAO was modeled theoretically using ChemBioDraw ultra and Marvin Sketch software that utilizes DFT (density functional theory) approach. Fig. 7 (a) showing the ground state configuration of TMAO as the space-filling model after optimization as well as energy minimization. The energy value of optimized molecule was found to be E(RB3LYP) = - 249.68522468 a.u., while RMS Gradient Norm = 0.00830651 a.u. with dipole moment = 4.6272 Debye. The greyish color balls represent C atoms, white color ball hydrogen atoms while ball with blue color symbolizes nitrogen atom and pink color represent oxygen atom of TMAO molecule.

**Fig. 7:**
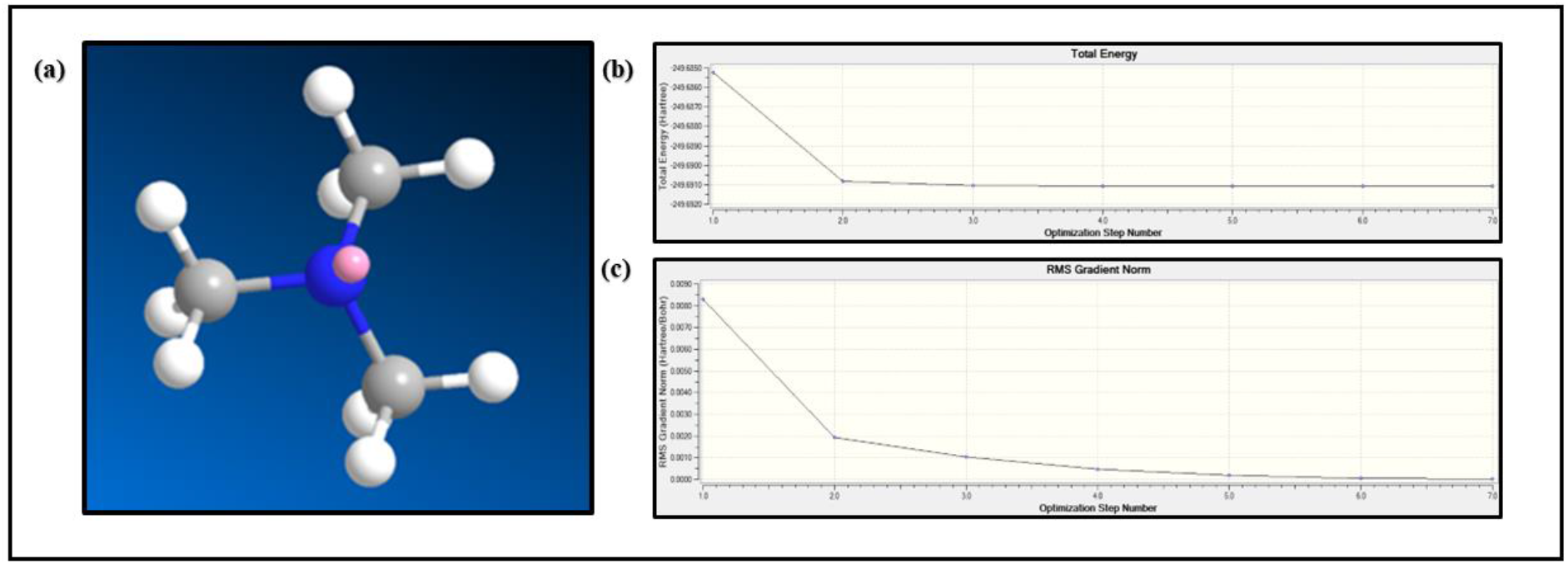
(a) 3D structure of ground state configuration of TMAO molecule, (b) Energy minimization and optimization graph of TMAO showing energy minimization with optimization step number (c) Energy minimization and optimization graph of TMAO showing energy RMS gradient normalization with optimization step number.

The different bond lengths (in Å) and bond angles (°) for all possible confirmations of atoms present in TMAO molecules in 3D space has been mentioned in detail in table S1 and S2 respectively.

#### 3.7.2 Docking of TMAO with BSA

HIS 67 and GLU 243 were observed to be involved in conventional hydrogen bond formation with TMAO. GLU 243, ASP 248 and GLU 251 are involved in electrostatic interaction while LYS 242 is involved in carbon hydrogen bond as shown in Fig.8.

**Fig. 8:**
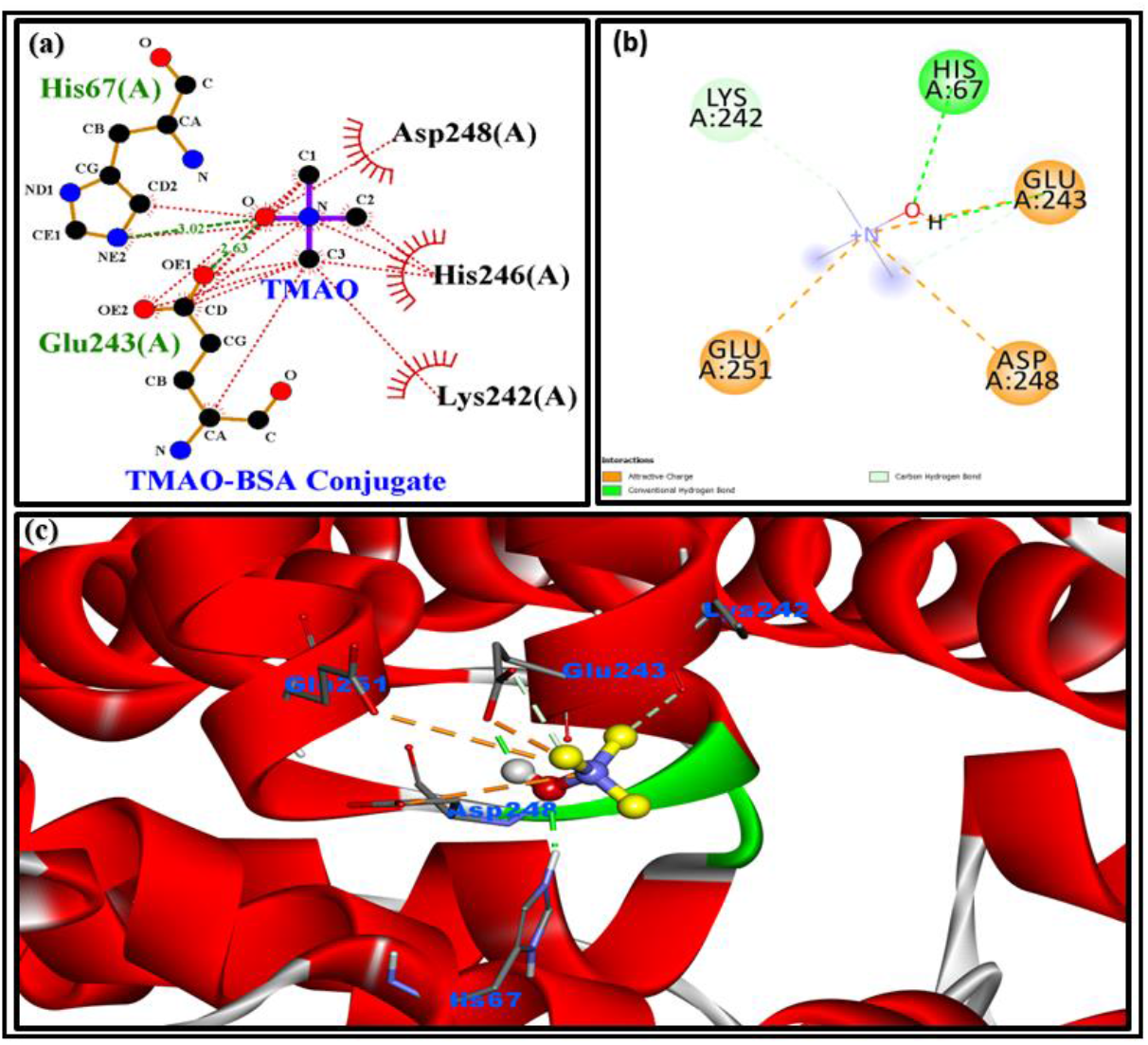
shows the TMAO-BSA docked complex, having involvement of electrostatics and hydrogen bonding of TMAO with BSA Fig.8: (a) shows the lig plot structure of interaction of different amino acid residues with different atoms of TMAO. (c) 2D interactions and (d) 3D interaction of TMAO-BSA conjugate.

From docking result, it is clearly shown that the TMAO is binding with BSA. Since the TMAO is very small molecule so interaction is weak here as evident from the binding energy score value is – 3.6 kcal/mol (shown in Table S3).

## 4. Conclusion

This manuscript describes about the interaction of TMAO with BSA with the help of various optical techniques and computational methods. Both the results obtained from experimental spectroscopic methods and *in silico* docking studies are in well agreement with each other. UV-Vis absorbance spectroscopy indicated the absorbance quenching of BSA with the increasing concentration of TMAO which could mainly be due to TMAO-BSA conjugate formation. The decrease in intensity of fluorescence is mainly due to involvement of tryptophan during the conjugation. The occurrence of static quenching was confirmed via Stern–Volmer (SV) plot, which consider the complex formation between fluorophore and protein molecule. The negative value for ΔG revealed that binding process is spontaneous in this case i.e., conjugate formation was occurred. From *insilco* analysis, we concluded the involvement of hydrogen bonding and electrostatic interaction are the major factor for stabilizing the TMAO-BSA complex. The link of TMAO with various diseases such as diabetes, renal and cardiovascular disease, thus aptamer against TMAO could be helpful in early diagnosis of the diseases related to it.

## Acknowledgments

Authors would highly acknowledge to Dr. Pratima Solanki and Dr. Anil Kumar to provide the experimental as well as computational facilities as well as entire proper coordinated guidance for research work.

## Contribution of Authors

AKV has performed entire research (experimental as well as computational work), analyzed all data and wrote the whole draft. PG has helped in experimental work and edited the draft. GBVS has helped in FTIR data analysis as well as reviewed the entire draft. PS and AK have designed the research, provided the experimental and computational facilities and reviewed the whole draft and also edited the paper for final version.

## Information of Author

