## Supplemental File for "Interaction studies of Gut metabolite; Trimethylene amine Oxide with Bovine Serum Albumin through Spectroscopic, DFT and Molecular Docking Approach"

The detail information for bond length, bond angle and dihedral angle for each atom of TMAO molecule has been mentioned below.

Table S1: Detail of Bond length (Å) for each atom of TMAO molecule

| | | C-H (CH3 Group) | C-H (CH3 Group) | C-H (CH3 Group) | N-O | N-C |
|---|---|---|---|---|---|---|
| | | **1.07** | **1.07** | **1.07** | | **1.47** |
| | **Unoptimized** | **1.07** | **1.07** | **1.07** | **1.36** | **1.47** |
| | | **1.07** | **1.07** | **1.07** | | **1.47** |
| **Bond length (Å)** | | | | | | |
| | | **1.08717** | **1.08716** | **1.08717** | **1.35851** | **1.50291** |
| | **Optimized** | **1.09395** | **1.09397** | **1.094** | | **1.50291** |
| | | **1.08715** | **1.08717** | **1.08716** | | **1.50288** |

Table S2: Detail of Bond angle (°) for each atom of TMAO molecule

| | | **H-C-H (CH3)** | **H-C-H (CH3)** | **H-C-H (CH3)** | **C-N-C** | **O-N-C** |
|---|---|---|---|---|---|---|
| | Unoptimized | **109.47122** | **109.47122** | **109.47122** | **109.47122** | **109.47122** |
| | | **109.47122** | **109.47122** | **109.47122** | **109.47122** | **109.47122** |
| | | **109.47122** | **109.47122** | **109.47122** | **109.47122** | **109.47122** |
| **Bond Angle (°)** | | | | | | |
| | | **111.25714** | **111.26552** | **111.25639** | **109.44178** | **109.50281** |
| | Optimized | **111.25799** | **111.26319** | **111.26253** | **109.44996** | **109.49708** |
| | | **109.09407** | **109.10266** | **109.11291** | **109.44996** | **109.4932** |

Table S3: Rank wise confirmation of TMAO-BSA complex with corresponding energy

| mode | affinity (kcal/mol) | dist from best mode rmsd l.b. | rmsd u.b. |
|---|---|---|---|
| 1 | -3.6 | 0.000 | 0.000 |
| 2 | -3.5 | 18.072 | 18.499 |
| 3 | -3.3 | 16.954 | 17.320 |
| 4 | -3.2 | 60.654 | 60.995 |
| 5 | -3.2 | 13.014 | 13.176 |
| 6 | -3.2 | 67.847 | 67.998 |
| 7 | -3.1 | 56.876 | 57.191 |
| 8 | -3.1 | 61.565 | 61.896 |
| 9 | -3.0 | 62.772 | 63.108 |

Writing output ... done.
